# Comparative proteome analysis of different *Saccharomyces cerevisiae* strains during growth on sucrose and glucose

**DOI:** 10.1101/2022.11.03.515096

**Authors:** Carla Inês Soares Rodrigues, Maxime den Ridder, Martin Pabst, Andreas K. Gombert, Sebastian Aljoscha Wahl

## Abstract

Both the identity and the amount of a carbon source present in laboratory or industrial cultivation media have major impacts on the growth and physiology of a microbial species. In the case of the yeast *Saccharomyces cerevisiae*, sucrose is arguably the most important sugar used in industrial biotechnology, whereas glucose is the most common carbon and energy source used in research, with many well-known and described regulatory effects, e.g. glucose repression. Here we compared the label-free proteomes of exponentially growing *S. cerevisiae* cells in a defined medium containing either sucrose or glucose as the sole carbon source. For this purpose, bioreactor cultivations were employed, and three different strains were investigated, namely: CEN.PK113-7D (a common laboratory strain), UFMG-CM-Y259 (a wild isolate), and JP1 (an industrial bioethanol strain). These strains present different physiologies during growth on sucrose; some of them reach higher specific growth rates on this carbon source, when compared to growth on glucose, whereas others display the opposite behavior. It was not possible to identify proteins that commonly presented either higher or lower levels during growth on sucrose, when compared to growth on glucose, considering the three strains investigated here, except for one protein, named Mnp1 – a mitochondrial ribosomal protein of the large subunit, which had higher levels on sucrose than on glucose, for all three strains. Interestingly, following a Gene Ontology overrepresentation and KEGG pathway enrichment analyses, an inverse pattern of enriched biological functions and pathways was observed for the strains CEN.PK113-7D and UFMG-CM-Y259, which is in line with the fact that whereas the CEN.PK113-7D strain grows faster on glucose than on sucrose, the opposite is observed for the UFMG-CM-Y259 strain.

## Introduction

The yeast *Saccharomyces cerevisiae* can utilize different carbohydrates as carbon sources. Transcription of genes involved in these processes is regulated according to nutrient availability, through signaling pathways that mainly respond to the levels of glucose in the environment ^1,2^. This yeast species prefers glucose over other sugars, which suggests that this substrate should lead to higher maximum specific growth rates (μ_MAX_) on glucose than on other carbon sources.

When *S. cerevisiae* is grown in batch cultivations using sucrose as the sole carbon source, some glucose and fructose will be present in the medium as a result of sucrose hydrolysis via periplasmic invertase, but the levels achieved will be lower, when compared to the ones during glucose batch conditions^3^. Because of this, differences are expected in central carbon metabolism and related metabolic pathways, which will eventually be reflected in altered physiologies. In our previous work ^3^, the μ_MAX_ of the *S. cerevisiae* laboratory strain CEN.PK113-7D was shown to be lower for sucrose-grown cells compared to glucose-grown cells. This is consistent with regulation exerted by glucose over the use of other carbon sources, whose consumption is hampered by a complex mechanism named the catabolite repression response (CRR). Other designations exist for this phenomenon, e.g. carbon catabolite repression or glucose repression ^1,4^. Interestingly, this trait was not observed for another *S. cerevisiae* strain we also investigated previously ^3,5^, namely UFMG-CM-Y259, which was isolated from the bark of trees in Brazil. The same situation was observed for a third strain, named JP1, which was isolated from a sugarcane biorefinery used to produce fuel ethanol in the Northeast region of Brazil ^6^. Both strains displayed higher μ_MAX_ values during growth on sucrose compared to glucose in batch bioreactor cultivations. Furthermore, in other published works reporting the use of different cultivation systems, a phenotype of faster growth on sucrose was also observed: in aerobic batch in microtiter plates ^5^, in batch bioreactor cultivations ^7^, or using a strain adapted on sucrose for 250 generations ^8^. This is especially surprising taking into account that sucrose is mainly degraded extracellularly by invertase into glucose and fructose, before these monosaccharides can be catabolized. Thus, this additional metabolic step requires the cells to synthesize and secret at least one additional protein, which is expected to represent an additional burden that could potentially lead to a decrease in the specific growth rate.

Faster growth is accompanied by an increased energy demand of the anabolic metabolism. Cells ensure that sufficient energy, in the form of adenosine triphosphate (ATP), is provided by modulating the flux through central carbon metabolism. The biochemical capacity of carbon metabolism, in turn, depends on the abundance of the metabolic enzymes in conjunction with their regulation ^9,10^. Many of these proteins undergo metabolite allosteric binding or post-translational modifications (PTM) that control their activity, such as the phosphorylation of phosphofructokinase 2 (Pfk2p) ^11^ and pyruvate kinase (Pyk1p) ^12^. Thus, protein abundance and the interactions among them strongly influence the cellular phenotype. Proteome measurements can reveal changes between conditions and facilitate the identification of cellular responses as well as elucidate regulatory mechanisms under diverse growth conditions and strains ^13–16^.

Known regulatory mechanisms include the protein kinase A (PKA) that plays a key role in the transcriptional regulation of genes encoding ribosomal proteins, thus being involved in ribosome biogenesis, as well as in stress responsive genes, such as *RIM15, MSN2* and *MSN4*. Moreover, this kinase also regulates the activity of metabolic enzymes at the post-translational level during carbon source shifts ^2,17^. The latter includes regulation of proteins for the synthesis and degradation of storage carbohydrates ^18–20^, for the glycolytic pathway ^11,21^, and for gluconeogenesis ^22^. Specifically for the glycolytic enzymes, PKA directly phosphorylates and activates the proteins Pfk2p and Pyk1p, which implies a positive correlation between PKA activity and the glycolytic flux. Both substrates, sucrose and glucose, induce the signaling cascade leading to PKA activation via the Gpr1p-Gpa2p protein coupled receptor ^23^. This receptor has a higher affinity for the disaccharide ^23^, leading to the hypothesis that PKA is involved in enabling a higher growth rate on sucrose ^24^.

To verify how regulatory mechanisms might vary among different strains of *S. cerevisiae*, we compared the proteomes of *S. cerevisiae* strains CEN.PK113-7D, JP1, and UFMG-CM-Y259 during growth on sucrose and the corresponding proteomes during growth on glucose. We performed comparative proteomic analysis using label-free quantification of protein abundance via mass spectrometry. Statistically significant changes in the proteome profiles related to glucose conditions were subjected to Gene Ontology (GO) overrepresentation and KEGG pathway enrichment analyses.

## Material and Methods

### Yeast strains and cultivation

Three *Saccharomyces cerevisiae* strains were used in this work: the laboratory strain CEN.PK113-7D ^25^, the strain UFMG-CM-Y259, which was isolated from a Brazilian tree ^5^, and the bioethanol strain JP1 ^26^. Yeast strains were pre-cultured in shake flask (Certomat BS-1, Braun Biotech International) with synthetic medium ^27^ containing urea as nitrogen source (2.3 g l^-1^) and supplemented with either 1% (w/w) glucose or the hexose equivalent amount of sucrose. After 24 h of growth at 30 °C at 200 rpm, a 1-ml aliquot of this pre-culture was transferred to another flask containing fresh preculture medium. The cells were cultured for 24 h, then harvested by centrifugation (3500 *g)* and washed twice with fresh cultivation medium. The bioreactor cultivation medium was prepared according to ^27^ with 2% glucose or the hexose equivalent amount of sucrose as sole carbon and energy source. The batch bioreactor cultivation with a working volume of 1.2 l (2 l vessel, Applikon Biotechnology B.V., Delft, The Netherlands) was started with an optical density of 0.2 at 600 nm by inoculation with cells resuspended in fresh cultivation medium. Aerobic bioreactor cultivations were carried out at 30 °C, 800 rpm, and with 0.5 l.min^-1^ air flow rate using a mass flow controller (Model 58505, Brooks Instrument, Hatfield, USA). The dissolved oxygen was above 40% throughout the whole cultivation for all conditions. The culture pH was kept at 5.0 by automatic addition of 0.5 mol.l^-1^ KOH (BioStat controller, type 8843415, Sartorius AG, Germany).

### Sample preparation, protein extraction and trypsin proteolytic digestion

Samples for proteomic analysis were collected during the exponential growth phase (EGP). Aliquots with approximately 2-mg protein content were withdrawn from the bioreactor cultivations and immediately centrifuged at 867 *g* for 5 minutes. The supernatant was discarded and the pellet frozen at −80 °C until further use. Cell pellets were resuspended in 100 mM TEAB lysis buffer containing 1% SDS and phosphatase/protease inhibitors. Protein extraction was performed using glass bead milling (425 – 600 μm, acid-washed) through a 10-time shaking for 1 min with a bead beater alternated with 1 min rest on ice. Proteins were reduced by addition of 5 mM DTT and incubated for 1 h at 37 °C. Subsequently, the proteins were alkylated for 60 min at room temperature in the dark by addition of 15 mM iodoacetamide. Protein precipitation was performed adding four volumes of ice-cold acetone (−20 °C) and proceeded for 1 h at −20 °C. The proteins were solubilized using 100 mM ammonium bicarbonate. Proteolytic digestion was performed by Trypsin (Promega, Madison, WI), 1:100 enzyme to protein ratio, and incubated at 37° C overnight. Solid phase extraction was carried out with an Oasis HLB 96-well μElution plate (Waters, Milford, USA) to desalt the mixture. Eluates were dried using a SpeedVac vacuum concentrator at 45° C. Dried peptides were resuspended in 3% ACN/0.01% TFA prior to MS-analysis to give an approximate concentration of 250 ng per μl. All chemicals used for sample preparation were supplied by Sigma-Aldrich (Missouri, USA).

### Label-free shot-gun proteomics

An aliquot corresponding to approx. 250 ng protein digest was analysed using an one dimensional shot-gun proteomics approach ^28^. Briefly, the samples were analysed using a nano-liquid-chromatography system consisting of an ESAY nano LC 1200 (Thermo Scientific, Waltham, USA), equipped with an Acclaim PepMap RSLC RP C18 separation column (50 μm x 150 mm, 2μm) (Thermo Scientific, Waltham, USA), and an QE plus Orbitrap mass spectrometer (Thermo Scientific, Waltham, USA). The flow rate was maintained at 350 nl.min^-1^ over a linear gradient from 6% to 26% solvent B over 65 minutes, followed by an increase to 50% solvent B over 20 min and a subsequent back equilibration to starting conditions. Data were acquired from 2.5 to 90 min. Solvent A was H2O containing 0.1% formic acid, and solvent B consisted of 80% acetonitrile in H2O and 0.1% formic acid. The Orbitrap was operated in data dependent acquisition mode acquiring peptide signals from 385-1250 m/z at 70K resolution. The top 10 signals were isolated at a window of 2.0 m/z and fragmented using a NCE of 28. Fragments were acquired at 17.5K resolution.

### Data analysis

Raw data were mapped to the proteome database from *Saccharomyces cerevisiae* (Uniprot, strain ATCC 204508 / S288C, Tax ID: 559292, July 2018) using PEAKS Studio X (Bioinformatics Solutions Inc, Waterloo, Canada) ^29^ allowing for 20 ppm parent ion and 0.02 m/z fragment ion mass error, 2 missed cleavages, carbamidomethyl as fixed and methionine oxidation and N/Q deamidation as variable modifications. To limit false-positive peptide identification, 1% false discovery rate (FDR) was applied to peptide spectrum matches (PSM), and subsequently the identified peptides were filtered with ≥ 2 unique peptides in all three biological replicates. Label free quantitative analysis was carried out on the identified peptides by using such module in the PEAKS Q software tool (Bioinformatics Solutions Inc, Waterloo, Canada) ^29^. The peak areas were normalized to the total ion count (TIC) of the respective analysis run before performing pairwise comparison between the carbon sources for each strain. Protein abundance ratio between sucrose and glucose conditions was filtered with fold change ratios ≥ 1.2, and analysis of variance (ANOVA) was performed to test the statistical significance of the observed abundance changes, with *p* values below 0.05 considered to be statistically significant.

### Gene Ontology (GO) term overrepresentation and pathway enrichment analyses of differentially expressed proteins

Differentially abundant proteins for each sucrose-glucose comparison were subjected to GO overrepresentation ^30^ as well as Kyoto Encyclopedia of Genes and Genomes (KEGG) ^31^ enrichment analyses. GO overrepresentation analysis was carried out using PANTHER ^32^ directly through the GO webpage (http://geneontology.org) and visualized applying the ggplot2 package (R 4.0.0). For this, GO biological processes were obtained with all identified proteins used as background reference. Fisher’s exact test with FDR multiple test correction were set for the GO overrepresentation analysis ^14^. KEGG enrichment analysis was performed using the *clusterProfiler* ^33^ (version 3.16.0) package within the Bioconductor project ^34^ in the R environment. Data visualizations were performed using the ggplot2 package in the R environment.

## Results

### Protein identification and differential expression on sucrose relative to glucose conditions

We identified a total of 237,916 peptides across all six conditions performed (triplicate injections), of which 30,197 were unique, resulting in 2,311 proteins identified

Changes in the proteome between the two growth conditions were identified for each strain by calculating the protein abundance fold changes (FC) with respect to the glucose growth condition. The significance of each observed fold change was determined by calculating *p* values. A FC >= 1.2 and an adjusted *p*-value <= 0.05 were defined as the biological and statistical threshold values, respectively. The comparative analysis revealed that there is no common pattern of sucrose-induced changes of protein levels amid all three studied strains. Nevertheless, there were pairwise overlaps (**Figure 1A, B**). For the CEN.PK113-7D strain, the abundance of 418 proteins were found statistically significant different (272 up; 146 down), against 510 (242 up; 268 down) and 224 (183 up; 41 down) proteins for the UFMG-CM-Y259 and JP1 strains, respectively. Remarkably, a cross strain analysis showed only one common protein with increased concentration on sucrose: Mnp1p – a mitochondrial ribosomal protein of the large subunit. No commonly decreased protein levels were observed for all three strains using the defined criteria.

**Figure 1.**
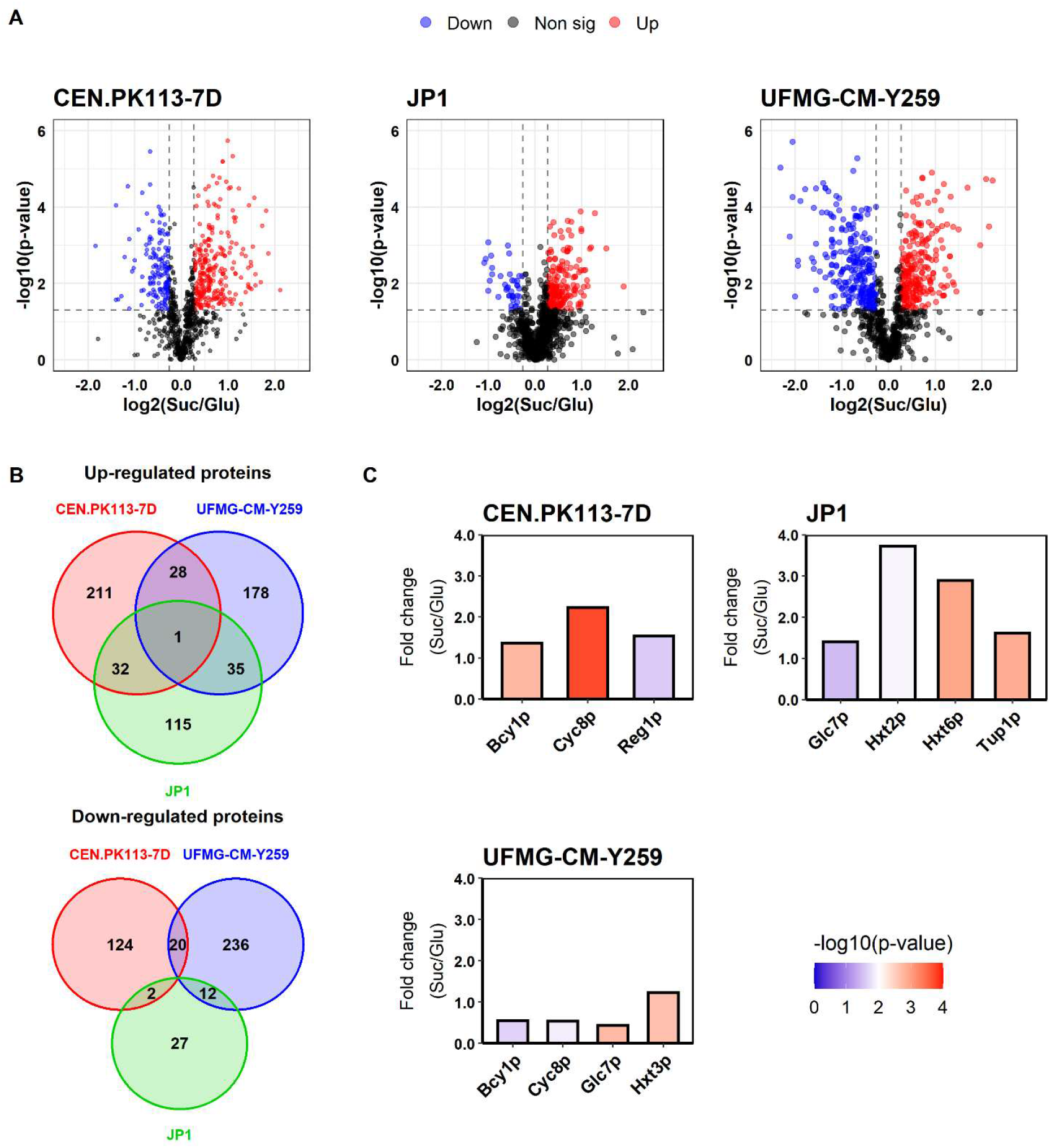
Comparative protein abundance between sucrose and glucose growth conditions for *Saccharomyces cerevisiae* CEN.PK113-7D, UFMG-CM-Y259, and JP1 strains. (A) Volcano plots highlight proteins with statistically significant differential abundance between growth on either substrate as sole carbon and energy source, considering a significance level of *p* < 0.05. Increased protein levels on sucrose over the glucose condition are shown in red, decreased proteins levels in blue and grey otherwise. (B) Venn diagrams illustrate the extent of differentially abundant proteins shared among the strains. (C) Fold changes in the levels of some proteins known to play a role in sucrose metabolism and regulation, hexose transport, and in the protein kinase A (PKA) signaling cascade. Color coding indicates the statistical significance.

Considering pair-wise comparisons, the following observations can be made: The CEN.PK113-7D strain shared 29 and 33 proteins that had increased levels on sucrose with the UFMG-CM-Y259 and JP1 strains. Furthermore, it shared 20 and 2 with decreased levels on sucrose with the same strains (see Fig. 1B). Between UFMG-CM-Y259 and JP1 strains, 36 proteins were mutually increased, whereas 12 were jointly decreased. It is important to highlight, that there was no transcription factor (TF) among the common proteins. However, individually, the UFMG-CM-Y259 strain showed increased abundance of the TF Rtg2p, and decreased abundance of the TFs Cyc8p, Hmo1p, and Wtm1p when comparing sucrose to glucose conditions. In the CEN.PK113-7D strain, in turn, the TFs Cyc8p, Wtm1p, and Rap1p had increased levels, whereas Hog1p presented decreased levels in the sucrose condition (**Supplementary material**).

#### Sucrose metabolism related proteins

With no need for sucrose hydrolysis during growth on glucose, we especially expected different levels for the protein invertase between the two growth conditions. Furthermore, the transcription of the *SUC2* gene, which is the most common allele encoding for this enzyme, is subject to glucose repression. However, invertase abundance was not detected as significantly changed for the UFMG-CM-Y259 and JP1 strains with the chosen criteria (**Figure 1C**). This enzyme was not identified in samples originating from the CEN.PK113-7D strain, suggesting an abundance below detection limit. This is in agreement with the low invertase activity observed earlier for the CEN.PK113-7D strain cultivated on sucrose ^3^.

Notably, for all strains, altered abundance in at least one of the proteins known to be involved in the SUC2regulation ^1,35^ was observed (**Figure 1C**). The general transcription co-repressor Cyc8p was significantly changed in both CEN.PK113-7D and UFMG-CM-Y259 strains, although in different manners: Increased in CEN.PK113-7D and decreased in UFMG-CM-Y259. For the JP1 strain increased levels of Tup1p, which is a general repressor of transcription that forms a complex with Cyc8p, were found on sucrose. This strain also exhibited increased levels of the protein phosphatase catalytic subunit Glc7p in the sucrose condition, contrary to the observations for UFMG-CM-Y259. Besides, Glc7p’s regulatory subunit, Reg1p, had increased levels only in the CEN.PK113-7D strain. The increased levels of both regulatory proteins Glc7p and Tup1p during the cultivations with sucrose for the JP1 strain suggest repression of the *SUC2* gene, although it is required during cultivations with the disaccharide. It has to be noted that the glucose concentration in the medium was above the reported threshold concentrations of 0.5, 2-3.2 g.l^-1^, reported for *SUC2* repression ^36–38^ at the sampling time points during the sucrose experiments with both UFMG-CM-Y259 and JP1.

#### Protein kinase A

Protein kinase A (PKA) activity has been reported to influence protein expression in response to the available carbon sources ^2,17^. Especially, PKA influences the transcription of growth-related genes and key glycolytic enzymes. Therefore, a correlation between growth rate and the activity of this kinase was expected.

For the CEN.PK113-7D strain, a 30% decrease (FC of 0.68) in the maximum specific growth rate (μ_MAX_) was observed, when comparing sucrose to glucose conditions ^3^. The proteome measurements described here, show increased levels for PKA’s regulatory subunit Byc1p in the CEN.PK113-7D strain grown on sucrose (FC = 1.37, **Figure 1C**). The increased levels could imply a reduced PKA activity, since Byc1p inhibits PKA’s catalytic subunits Tpk1/2/3p. In contrast to the CEN.PK113-7D strain, Byc1p levels were reduced in UFMG-CM-Y259 (FC = 0.55, Fig. 1C). For this strain, a 28% increase in μ_MAX_ was reported for growth on sucrose ^3^, suggesting enhanced PKA activity for the sucrose condition. No significant different level of Bcy1p was detected in the JP1 strain. The redundant catalytic subunits of PKA could not be quantified in any of the three strains.

The activity of PKA is mainly regulated by the binding of cyclic adenosine monophosphate (cAMP) to its regulatory subunit, through a signaling pathway mediated by the RAS proteins which is triggered by the presence of glucose or sucrose sensed by the G protein coupled receptor ^39,40^. Strain CEN.PK113-7D has a mutation in the adenylate cyclase encoding gene *CYR1*, resulting in a modified phenotype in terms of storage metabolism and cAMP was not detected in the metabolome ^41^. This could explain why sucrose did not positively affect PKA activity in CEN.PK113-7D cells, despite the higher affinity of the *GPR1* receptor for sucrose compared to glucose. Alternatively to cAMP binding, Kelch-repeat proteins (Krh1/2p) can control PKA activation ^42,43^. However, CEN.PK113-7D cells are also unable to produce these proteins ^41^. With the current literature knowledge, the mechanism by which this strain regulates PKA activity remains unknown.

#### Hexose transporters

As the expression of hexose transporters in *S. cerevisiae* is sensitive to the extracellular concentrations of hexoses ^44,45^, we expected different levels for these proteins under glucose and sucrose conditions. For the JP1 strain, the low-affinity hexose transporters Hxt2p and Hxt6p were found at higher levels, whereas the UFMG-CM-Y259 strain showed higher Hxt3p levels (**Figure 1C**). For the CEN.PK113-7D strain, no change in hexose transporter levels could be quantified due to weak identification. This was also the case for other hexose transporters that were not quantified in JP1 and UFMG-CM-Y259.

The increased levels of Hxt3p in sucrose-grown UFMG-CM-Y259 cells is in agreement with a previous report using wine yeast, claiming that this protein is involved in enhanced fructose utilization ^46^. However, the unchanged level of this protein in the JP1 strain suggests its regulation is strainspecific.

### Gene Ontology term overrepresentation and pathway enrichment analyses of differently expressed proteins

Next to the targeted analysis of differentially expressed proteins, gene ontology (GO) term overrepresentation and pathway enrichment analyses were performed to identify changes in biological functions. These analyses were based on pairwise comparisons, separately for increased and decreased protein levels.

For the JP1 strain, neither a significantly overrepresented biological process nor any enriched pathways ^3^ could be observed, comparing growth on sucrose and glucose. Therefore, in the following analysis, strain JP1 will not be included.

The GO enrichment analysis applied for the CEN.PK113-7D strain showed that proteins with increased levels on sucrose were mainly associated with translation, and downstream protein processes (protein metabolic process, cellular protein metabolic process). Proteins that decreased were involved in organic acid metabolic process and amino acid metabolic processes (**Figure 2**). For the UFMG-CM-Y259 strain a reverse pattern of represented biological processes to that of CEN.PK113-7D was observed: Biosynthetic process proteins increased during sucrose growth, while proteins associated to translation decreased. A massive increase was observed for ‘Metabolic Process’ containing a large set of catabolic and biosynthetic genes. This category was not found different for CEN.PK-113-7D.

**Figure 2.**
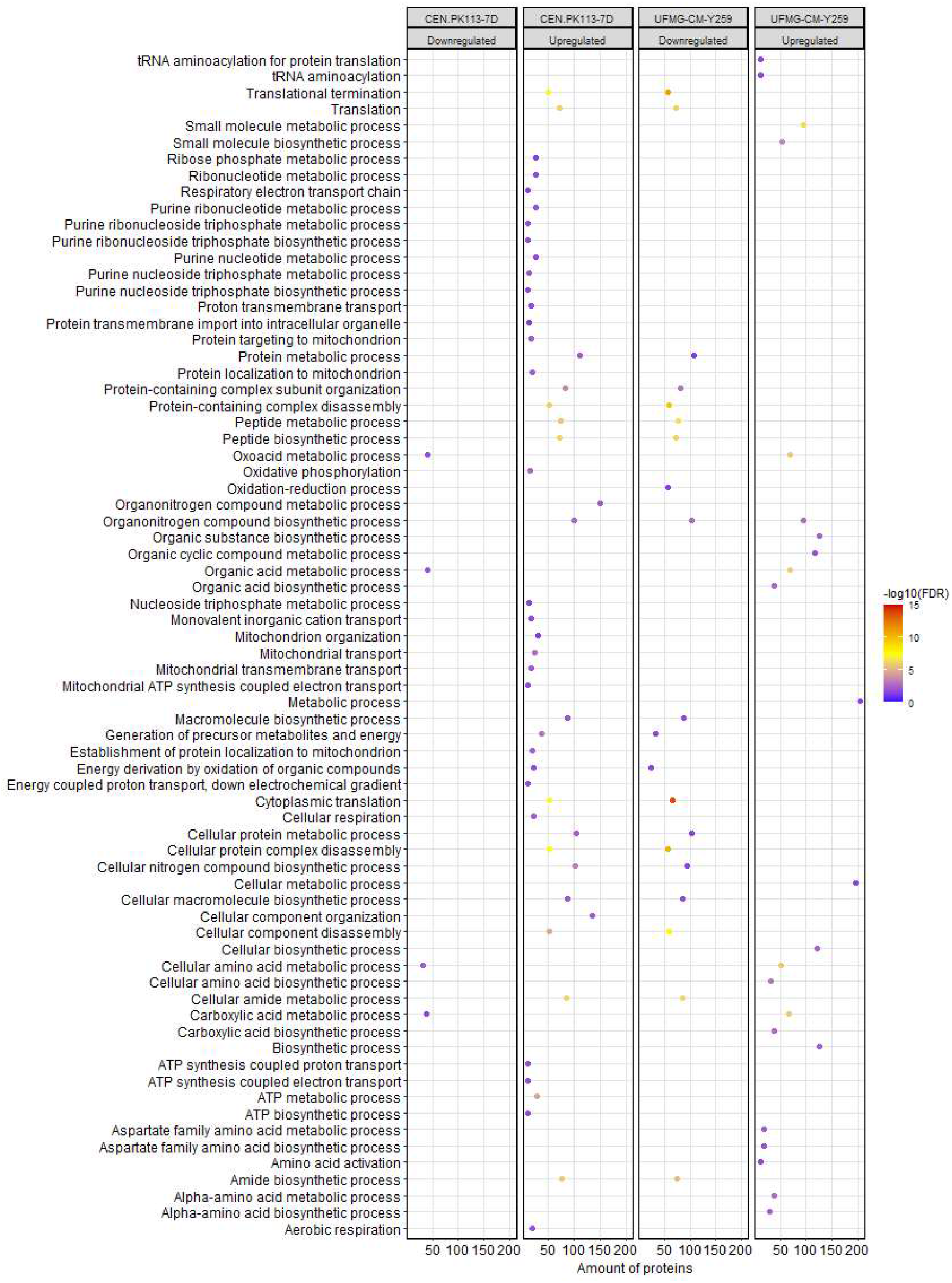
GO biological process overrepresentation analysis for differentially expressed proteins for *Saccharomyces cerevisiae* strains CEN.PK113-7D and UFMG-CM-Y259 cultivated on sucrose compared to glucose grown cells. The number of proteins in the dataset that falls in each overrepresented biological process is shown on the x-axis (amount of proteins).

KEGG pathway enrichment analysis showed similar results. For CEN.PK113-7D, proteins related to ribosome and respiratory metabolism increased under sucrose conditions (**Figure 3**). A decrease was observed for protein levels that play a role in biosynthesis of amino acids as well as of secondary metabolites. Similar to the GO biological enrichment analysis, for the UFMG-CM-Y259 strain, the proteins which decreased levels can be found in the ribosome pathway. Moreover, KEGG annotation identifies proteins in both increased and decreased groups with function in the biosynthesis of secondary metabolites, although with a higher relative proportion (GeneRatio) for the decreased protein levels.

**Figure 3.**
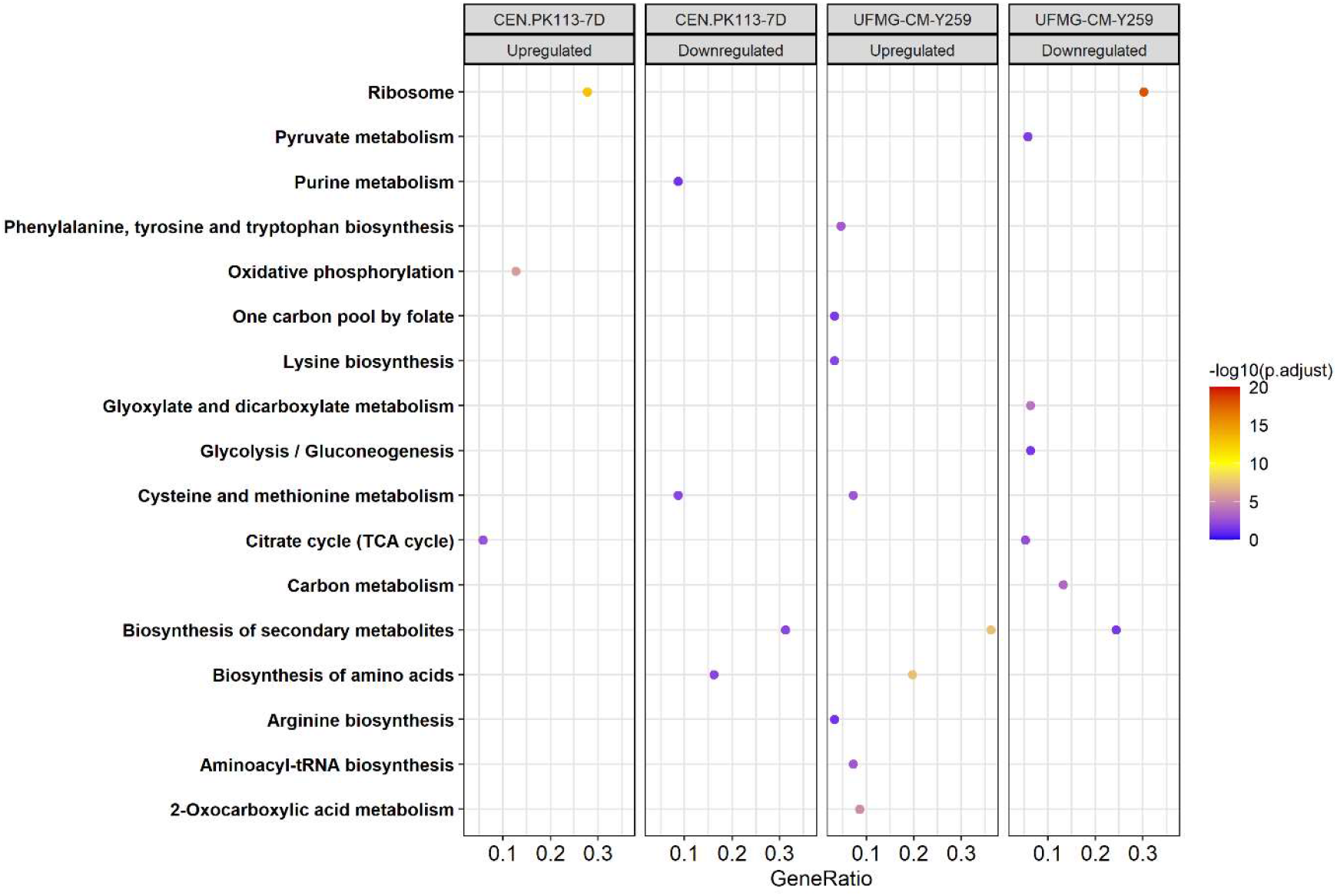
Pathway enrichment analysis for statistically significant proteins with an adjusted p-value < 0.05 for *S. cerevisiae* strains CEN.PK113-7D and UFMG-CM-Y259 grown on sucrose compared to glucose grown cells.
KEGG annotation was used for mapping the pathways. GeneRatio represents the proportion of proteins (gene products) in the correspondent abundance groups that is found in each category.

### Link between physiology and changes in protein abundance

For oxidative phosphorylation, as evidenced by pathway enrichment analysis, only the CEN-PK113-7D strain displayed a significant increase, which correlates with a 1.21 fold increase in the respiration rate (Sucrose: q_O2_ = 2.96 mmol/gDM/h, on glucose q_O2_ = 2.45 mmol/gDM/h) ^3^. For the UFMG-CM-Y259 strain a 1.27 fold increase in the specific oxygen uptake rate was observed for sucrose ^3^ (q_O2_ = 3.34 mmol/gDM/h, on glucose q_O2_ = 2.63 mmol/gDM/h, Table 1). Nevertheless, this higher q_O2_ was not accompanied by increased abundance of cytochrome c oxidase, except for its subunit 13. Furthermore, several proteins involved in the ergosterol and heme biosynthesis had increased (**supplementary Table S1**). Taking these observations into account, there seems to be a higher demand for molecular oxygen in the UFMG-CM-Y259 strain to sustain non-respiratory pathways that require O_2_ ^47^ during faster growth on sucrose. For the industrial strain JP1, no significant changes were found in either the oxygen uptake rate (q_O2_ = 2.19 mmol/gDM/h on sucrose and q_O2_ = 2.12 mmol/gDM/h on glucose), nor in protein abundance.

**Table 1.** **Comparison of metabolic rates and respective protein level** of *S. cerevisiae* CEN.PK113-7D, UFMG-CM-Y259 and JP1 during growth on sucrose relative to growth on glucose as sole carbon and energy source. Rates are taken from ^3^.

|  | Fold Change |  |  |
| --- | --- | --- | --- |
|  | CEN.PK113-7D | UFMG-CM-Y259 | JP1 |
| **Metabolic rates and yields** |  |  |  |
| qO<sub>2MAX</sub> | 1.21 | 1.27 | 1.03 |
| Molar fraction of substrate metabolized via the fermentative pathway | 0.87 | 1.10 | 0.97 |
| qS<sub>MAX</sub> | 0.62 | 1.07 | 1.05 |
| **Protein levels** |  |  |  |
| Oxidative phosphorylation | Up | N.S. | N.S. |
| Glycolysis/Gluconeogenesis | N.S. | Down* | N.S. |

N.S. = no significant change.
\* Proteins which reduced abundance: Acs1p, Adc1p, Ald4p, Eno1p, Lat1p, Lpd1p, Pck1p, Pda1p, Pfk2p, Pgm2p, Tdh2p.

Looking into catabolic rates (q_Smax_, as proxy for the glycolytic fluxes), minor differences were observed for either the UFMG-CM-Y259 (q_S_ = 12.84 mmol_GLCeq_/g_DM_/h on sucrose and q_S_ = 11.96mmol_GLCeq_/g_DM_/h on glucose) or the JP1 strain (q_S_ = 12.29 mmolGLCeq/gDM/h on sucrose and q_S_ = 11.67 mmolGLCeq/gDM/h on glucose), while for the CEN.PK113-7D strain a decrease in the maximum substrate specific consumption rate was observed (q_S_ = 8.23 mmolGLCeq/gDM/h on sucrose and q_S_ = 13.22 mmol_GLCeq_/g_DM_/h on glucose) (**Table 1**). Nevertheless, in the CEN-PK113-7D strain, the glycolytic enzyme levels were not affected by the change in substrate (**Figure 3; Table 1**). This suggests that the glycolytic flux is mainly controlled at the post-translational level, either by allosteric regulation of the glycolytic enzymes or by post transcriptional modifications, which is consistent with previous findings using continuous culture of the strain CEN.PK113-7D ^10,48^. For the JP1 and UFMG-CM-Y259 strains with increased glycolytic rates, no significant increase in respective protein abundance was observed, for UFMG-CM-Y259 even a small decrease in some enzyme concentrations was found (Table 1). Thus, there seems to be no correlation between protein level and obtained metabolic flux, a decoupling observed earlier for glycolysis. As discussed in an earlier section, the glycolytic protein levels have been reported to depend on the extracellular glucose concentration. The different strains and conditions lead to various situations that will be discussed in the following.

### Impact of the extracellular glucose concentration

The extracellular glucose concentration in each sucrose cultivation varied depending on the strain (**Supplementary Figure S1**). A high glucose concentration is known to repress the transcription of several genes associated to respiratory metabolism ^1^. The CEN.PK113-7D strain, was the one that displayed the lowest hexose concentrations in the extracellular environment during growth on sucrose (**Supplementary Figure S1**). The residual glucose concentrations observed in the medium for the UFMG-CM-Y259 and JP1 strains, during growth on sucrose, were one order of magnitude higher.

Consistent with the mentioned regulatory mechanism, proteins related to cellular aerobic respiration proteins had increased abundances in sucrose-grown CEN.PK113-7D cells (**Figures 2 and 3**). This resulted in a larger fraction of the substrate being metabolized via the respiratory pathway by this strain ^3^.

For the JP1 strain, the pathway analysis showed no overrepresented biological processes or enhanced pathways at either regulation level, suggesting that glucose repression was present for both substrate conditions. Furthermore, the use of fermentative metabolism by this strain was comparable under both substrate conditions (**Table 1**). For the UFMG-CM-Y259 strain slightly lower glucose concentrations were observed during sucrose cultivations, compared to JP1. Nevertheless, a small decrease in the enzyme levels of the tricarboxylic acid (TCA) was observed (**Figure 3**).

It has been described that the expression of TCA cycle proteins is glucose repressible under *HAP* control, especially when cells are cultivated on nitrogen sources that require α-ketoglutarate/glutamate for assimilation, such as ammonium used in this work. Furthermore, transcription of genes encoding for some proteins involved in the TCA cycle are influenced via the retrograde proteins Rtg1-3p ^17,49–51^. Here, the levels of the products of the retrograde target genes *CIT2, ACO1, IDP2*, and *IDH1/2* were decreased in the sucrose condition (**supplementary Table S1**). Especially, the TF Rtg2p level had an increased level during growth on sucrose suggesting that the retrograde pathway plays a role in controlling respiratory metabolism in the UFMG-CM-Y259 strain, parallel to the control exerted by the glucose repression phenomenon.

## Discussion

The comparison of protein levels of *S. cerevisiae* grown on sucrose compared to glucose showed specific changes in the different analyzed strains. No significant changes in abundance of transcription factors and proteins shared by all strains were observed, suggesting that no cellular function was under the exclusive control of sucrose.

With no obvious joint mechanisms, we correlated our findings with physiological data previously reported. The proteome measurements revealed a decrease of biosynthetic functions in favor of processes related to energy generation. In recent systems biology approaches, metabolic switches have been predicted and explained by resource allocation modeling ^52,53^. In short, complex pathways like respiration generate high stoichiometric amounts of ATP, but under protein space limitations become less favorable as the ATP/protein mass is lower compared to short, fermentative pathways. Therefore, Crabtree-positive yeasts, such as *S. cerevisiae*, switch to stoichiometrically less efficient fermentation under substrate excess conditions rather than oxidative phosphorylation, liberating proteome space. This space is filled with proteins for biomass production (biosynthesis of amino acids and secondary metabolites) allowing for a higher growth rate at lower yield. Note, that here several generations under batch conditions were monitored, the measured proteome can be expected to be a result of regulation. After long term evolution, the proteome allocation may shift.

The slower growth on sucrose relative to glucose observed for the CEN.PK113-7D strain could be a consequence of the regulatory mechanism not yet reaching the fastest growth for sucrose. For this strain lowest glucose levels were observed; The expression of respiration pathways is reported to be regulated via glucose repression which is in line with increased oxidative phosphorylation on sucrose. For other strains, especially the UFMG-CM-Y259 faster growth on sucrose over glucose was achieved with a reduced level of respiration and an increase in fermentation.

Additionally, we correlated protein kinase A to phenotypes of faster growth on sucrose than on glucose, which has been hypothesized earlier ^24^. Here, a positive relationship between PKA levels and maximum specific growth rate were found. However, as PKA functions via the post-translational modification mechanism of phosphorylation, an in-depth analysis of its contribution requires a different approach. Alternatively, phosphoproteomics could aid in untangling PKA and other signaling pathways to allow for a better understanding of the relationship between cellular growth and metabolism.

## Author contributions (names must be given as initials)

Everyone contributed to the experimental design. CIRS ran the cultivations. CIRS, MDR and MP ran the proteomic analyses. CIRS and MDR drafted the manuscript, which was further elaborated by AKG and SAW. Everyone read and approved the final version.

## Data availability statement (mandatory)

The MS/MS raw data is available via the PRIDE archive under accession number PXD037944

## Acknowledgements (optional)

We would like to acknowledge Fundação de Amparo à Pesquisa do Estado de São Paulo (FAPESP, São Paulo, Brazil) for funding and for scholarships to CISR (grant numbers 2016/07285-9, 2017/08464-7 and 2017/18206-5). This work was carried out as part of a Dual Degree Ph.D. project under the agreement between UNICAMP and Delft University of Technology.

