## Supplementary Material for "Comparative proteome analysis of different *Saccharomyces cerevisiae* strains during growth on sucrose and glucose"

**Table S1**. Summary of label free quantification and significance analyses for transcription factors, and proteins involved in retrograde response and ergosterol or heme biosynthesis for *S. cerevisiae* CEN.PK113-7D and UFMG-CM-Y259 cultivated in aerobic batch bioreactors with either sucrose or glucose as sole carbon source.

| Accession | Gene code | Log2_(Suc/Glu) | Significance | Function |
|---|---|---|---|---|
| | | **Transcription factor** | | |
| **CEN.PK113-7D** | | | | |
| Q12363 | WTM1 | 0.7312 | 27.22 | Transcriptional modulator WTM1 |
| P11938 | RAP1 | 0.6229 | 20.47 | DNA-binding protein RAP1 |
| P14922 | CYC8 | 1.1570 | 37.27 | General transcriptional corepressor CYC8 |
| P32485 | HOG1 | -0.4540 | 33.07 | Mitogen-activated protein kinase HOG1 |
| **UFMG-CM-Y259** | | | | |
| P32608 | RGT2 | 0.5059 | 14.01 | Retrograde regulation protein 2 |
| P14922 | CYC8 | -0.8890 | 18.61 | General transcriptional corepressor CYC8 |
| Q03973 | HMO1 | -0.4344 | 22.78 | High mobility group protein 1 |
| Q12363 | WTM1 | -0.6439 | 25.40 | Transcriptional modulator WTM1 |
| | | **Retrograde target** | | |
| **CEN.PK113-7D** | | | | |
| P41939 | IDP2 | -0.4941 | 14.57 | Isocitrate dehydrogenase [NADP] cytoplasmic |
| P28834 | IDH1 | 0.5059 | 44.72 | Isocitrate dehydrogenase [NAD] subunit 1 mitochondrial |
| P28241 | IDH2 | 0.3561 | 17.86 | Isocitrate dehydrogenase [NAD] subunit 2 mitochondrial |
| **UFMG-CM-Y259** | | | | |
| P08679 | CIT2 | -0.8890 | 24.20 | Citrate synthase peroxisomal |
| P19414 | ACO1 | -0.4540 | 24.11 | Aconitate hydratase mitochondrial |
| P41939 | IDP2 | -0.4540 | 28.86 | Isocitrate dehydrogenase [NADP] cytoplasmic |
| P28834 | IDH1 | -0.4739 | 17.81 | Isocitrate dehydrogenase [NAD] subunit 1 mitochondrial |

| Accession | Gene code | Log2_(Suc/Glu) | Significance | Function |
| --- | --- | --- | --- | --- |
| P28241 | IDH2 | -0.6666 | 52.80 | Isocitrate dehydrogenase [NAD] subunit 2 mitochondrial |
| **Ergosterol or heme biosynthesis** | | | | |
| **CEN.PK113-7D** | | | | |
| P54839 | ERG13 | 0.7991 | 40.05 | Hydroxymethylglutaryl-CoA synthase |
| P07143 | CYT1 | 0.5656 | 25.95 | Cytochrome c1 heme protein mitochondrial |
| P11353 | HEM13 | 0.6229 | 21.05 | Oxygen-dependent coproporphyrinogen-III oxidase |
| P21147 | OLE1 | 1.2690 | 21.52 | Acyl-CoA desaturase 1 |
| **UFMG-CM-Y259** | | | | |
| P32476 | ERG1 | 0.4436 | 17.94 | Squalene monooxygenase |
| P54781 | ERG5 | 0.5460 | 28.03 | Cytochrome P450 61 |
| P10614 | ERG11 | 0.2750 | 24.54 | Lanosterol 14-alpha demethylase |
| P07277 | ERG12 | 1.3951 | 19.63 | Mevalonate kinase |
| P54839 | ERG13 | 0.5850 | 18.04 | Hydroxymethylglutaryl-CoA synthase |
| P53199 | ERG26 | 0.7137 | 27.93 | Sterol-4-alpha-carboxylate 3-dehydrogenase decarboxylating |
| P32352 | ERG2 | -0.2863 | 17.05 | C-8 sterol isomerase |
| P32353 | ERG3 | -0.2863 | 15.41 | Delta(7)-sterol 5(6)-desaturase |
| P53045 | ERG25 | -2.0000 | 16.54 | Methylsterol monooxygenase |
| P09950 | HEM1 | 0.4647 | 19.40 | 5-aminolevulinate synthase mitochondrial |
| P05373 | HEM2 | 0.7398 | 17.85 | Delta-aminolevulinic acid dehydratase |
| P28789 | HEM3 | 1.0072 | 26.17 | Porphobilinogen deaminase |
| P40075 | SCS2 | 0.9411 | 21.38 | Vesicle-associated membrane protein-associated protein |

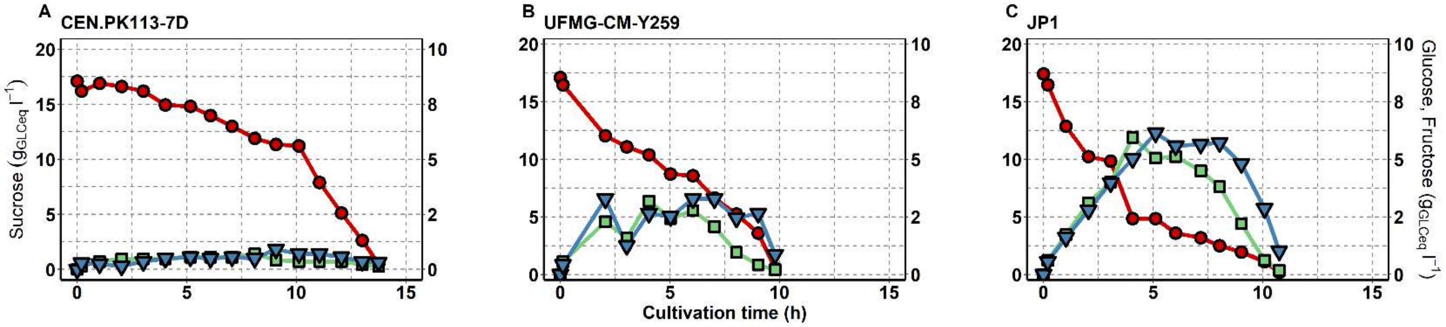


**Figure S1.** Sugar concentration during aerobic batch cultivations of *S. cerevisiae* CEN.PK113-7D, UFMG-CM-Y259, and JP1 in synthetic medium supplemented with sucrose as sole carbon and energy source.
